# Insights into the representativeness of biodiversity assessment in large reservoir through eDNA metabarcoding

**DOI:** 10.1101/2024.05.02.592166

**Authors:** Thainá Cortez, André LQ Torres, Murilo Guimarães, Henrique B Pinheiro, Marcelo Cabral, Gabriel Zielinsky, Camila M Pereira, Giovanni M de Castro, Luana TA Guerreiro, Juliana A Americo, Danielle LAS do Amaral, Mauro F Rebelo

## Abstract

Monitoring biodiversity on a large scale, such as in hydropower reservoirs, poses scientific challenges. Conventional methods such as passive fishing gear are prone to various biases, while the utilization of environmental DNA (eDNA) metabarcoding has been restricted. Most eDNA studies have primarily focused on replicating results from traditional methods, which themselves have limitations regarding representativeness and bias. In our study, we employed eDNA metabarcoding with three markers (12SrRNA, COI, and 16SrRNA) to evaluate the biodiversity of an 800 km² reservoir. We utilized hydrodynamic modeling to determine water flow velocity and the water renewal ratio throughout the study area. Additionally, we conducted statistical comparisons – rarefaction curves and multivariate methods – among samples as an alternative approach to assess biodiversity representation. The eDNA identified taxa previously documented in the reservoir by traditional monitoring methods, as well as revealed 29 – nine fishes and 20 non-fish – previously unreported species. These results highlight the robustness of eDNA as a biodiversity monitoring technique. Our findings also indicated that by randomly sampling 30% of the original number of samples, we could effectively capture the same biodiversity. This approach enabled us to comprehend the reservoir’s biodiversity profile and propose a straightforward, cost-effective monitoring protocol for the future based on eDNA.

## Introduction

Increasing the accuracy of biodiversity estimation has been a challenge in the face of lacking detailed information about population dynamics, habitat-based models, and either the occurrence or abundance of a species, especially for rare or elusive species [1–3]. Advancements in technology and methods have evolved over time from identifying individual organisms based on specific DNA barcodes to the current capability to analyze hundreds of thousands of DNA barcodes obtained from environmental samples (e.g. water, soil, air), allowing the description of entire communities in terms of species composition and dynamics [4]. This high-throughput approach, known as ‘DNA metabarcoding,’ is especially potent at uncovering biodiversity (e.g. 5–8).

Monitoring biodiversity through environmental DNA (eDNA) poses enormous challenges, especially in aquatic systems, due to the vast scales involved, and the combination of environmental conditions such as water current speed and density, wind, and tides, which influence particle transport [9]. Hydrodynamic modeling is an alternative approach for investigating the environmental influence on the transport of molecules. Such modeling approaches offer physical insights into the water body by considering outflow volume, water renewal rate, current velocity, and flow direction. This information together can enhance our understanding of the fundamental processes affecting the spatial and temporal (including seasonal) distribution of eDNA [10-12].

Hydroelectric Power Plant (HPP) reservoirs created by dam construction are constantly monitored, generating reports from long-term biodiversity assessments, which are crucial for tracking changes in aquatic communities. However, despite the importance of these long-term surveys, the validity of these longstanding monitoring techniques has been questioned by regulators and scientists, and their cost and utility questioned by operators [13]. The longstanding practice of the identification of species based on morphology relies upon the availability and knowledge of taxonomists, which is laborious, slow, and expensive, with high error rates [14]. In recent years eDNA has been gaining acceptance because it is unobtrusive and less costly [15, 16], yet highly effective [17]. Typically, eDNA metabarcoding studies are based on multiple sampling sites with several replicates at each location. Assuming that in aquatic systems eDNA is dispersed by water flow and begins to decay [18], higher numbers of sampling sites supposedly ensure the capture of the DNA of a wider diversity of species. Several approaches can help assess the optimal experimental design for a given study area. For instance, rarefaction curves can reveal the correlation between sampling size and the number of sequences per taxon (or Molecular Operational Taxonomic Unit, MOTUs), demonstrating the effectiveness of both the number of samples collected and the molecular approach [19].

Although this analysis is often employed in eDNA studies to determine whether the number of sampling sites or the molecular protocol sufficed to capture diversity [20–22], it has rarely been used to assess the optimal number of samples in large reservoirs [23, 24]. In this context, multivariate analyses, such as Principal Component Analysis (PCA) and Discriminant Analysis of Principal Components (DAPC), can offer insights into the distribution of variation across all sites, indicating which samples contribute most to the overall variability [19]. Finally, diversity analyses, including richness, Shannon’s H index, and species frequency computation, can complement these findings by providing visualizations that enable the recognition of community patterns and differences. These combined methods can inform how comprehensive the experimental design and the applied molecular approach were.

The aim of this study was to build a large-scale biodiversity inventory of a HPP reservoir utilizing eDNA and compare its findings to the previous reports either published in the scientific literature or filed over decades with regulators. The robustness of the inventory, as well as the most cost-effective experimental design and optimal number of samples, were investigated. Toward this goal, hydrodynamic modeling was initially conducted at an 800 km² HPP reservoir, Três Irmãos (TI, São Paulo, Brazil), coupled with pilot sampling at a single site to gain insight into local diversity using eDNA metabarcoding and three distinct markers (COI, 16SrRNA and 12SrRNA). Subsequently, multiple stations were sampled across the reservoir to evaluate large-scale biodiversity. Multivariate analyses, rarefaction curves, and community comparisons were employed to assess the effectiveness of the sampling design in compiling a comprehensive biodiversity inventory of the reservoir.

## Material and Methods

### Study area

The Três Irmãos (TI) hydroelectric power plant reservoir is located at the lower Tietê River basin, in São Paulo State in the Southeastern region of Brazil (Fig 1A). A 9,600 m long artificial channel called Pereira Barreto – the second-largest freshwater canal in the world – connects the TI HPP reservoir to the nearby Ilha Solteira HPP reservoir on the São José dos Dourados River (Fig 1A). This enables coordinated water management of the two reservoirs and integrated energy operation of the two hydroelectric plants, both of which drain into the Paraná River.

**Fig 1.**
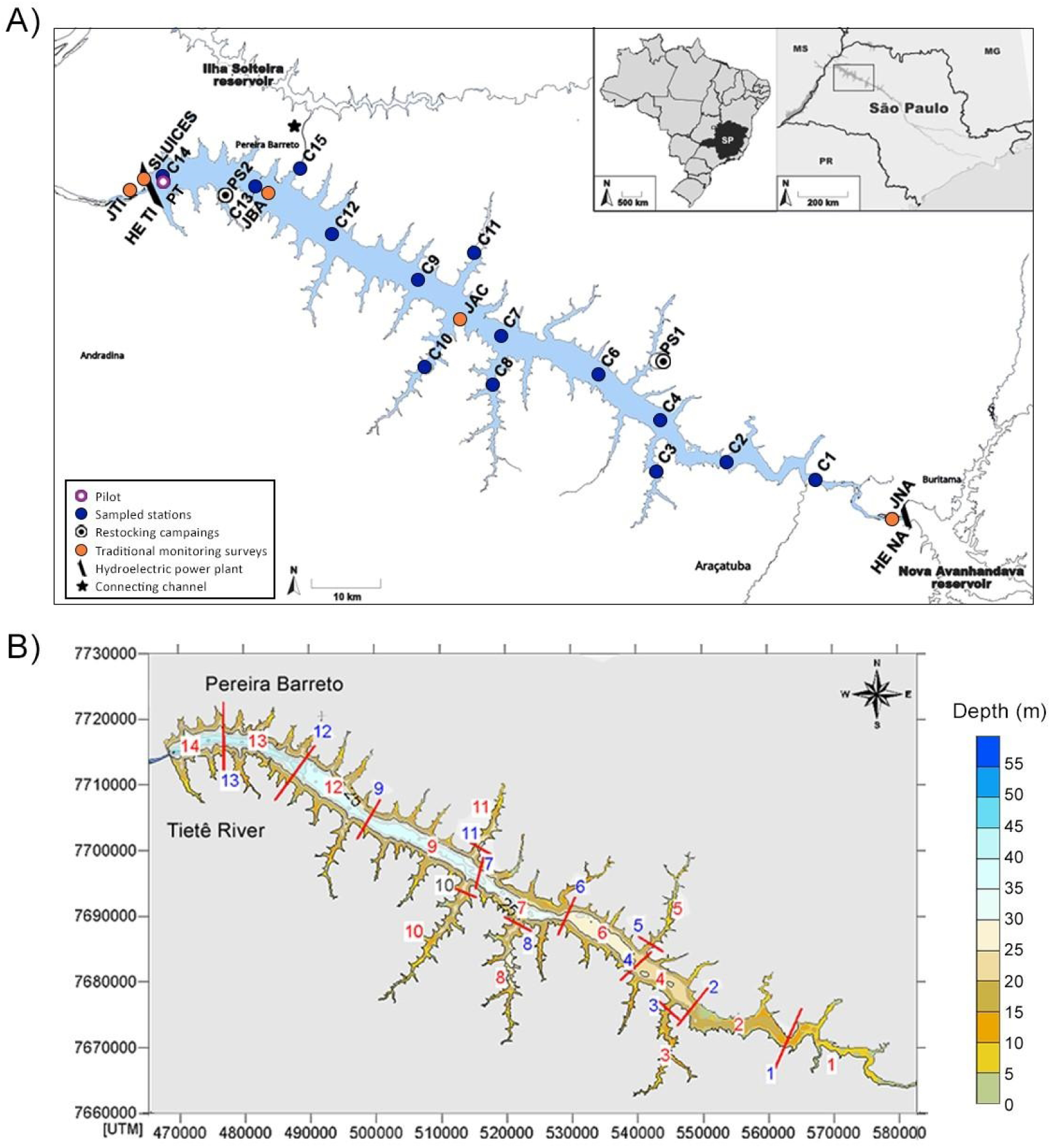
Três Irmãos (TI) reservoir and hydrodynamic modeling compartments. A) The TI and all sampling sites are shown. Purple circle shows the pilot sampling location. Blue circles labeled C1-C4 and C6-C15 designate the positions of the 14 sampling stations along the TI reservoir. Stations abbreviations as in S1 Table. White circles (PS1 and PS2) indicate where fish restocking programs are conducted. Orange circles (JNA, JAC, JBA, SLUICES and JTI) show the locations where the traditional taxonomic surveys are conducted by the hydroelectric company. Black trapezoid bars represent the positions of the Três Irmãos (UHE TI) and Nova Avanhandava (UHE NA) hydroelectric dams. Black star shows the Pereira Barreto service channel that connects the TI and IS reservoirs. B) For water renewal computations by hydrodynamic modeling, the reservoir was divided into 14 compartments. Red bars delineate the boundaries of the compartments. Compartment numbers are in red; the blue numbers are the control sections in which time series flows were calculated with the hydrodynamic model.

Três Irmãos is the largest power plant on the Tietê River. Its operation began in November 1993 (CESP 2013). The reservoir has a flooded area of 817 km^2^, a volume of 13,800 x 10^6^ m^3^, and an average depth of 17 m^3^. Inflow into the TI reservoir is limited primarily by the Nova Avanhandava (NA) HPP upstream of the reservoir, and secondarily by the flows across the Pereira Barreto channel which vary according to the operational requirements of the two hydroelectric power plants. The TI reservoir area has multiple uses, such as fish farming, professional and recreational fishery, tourism associated with boat trips, and cattle breeding, making it an anthropic area [25, 26].

### Hydrodynamic modeling

The modeling domain, used for hydrodynamic modeling, covered the entire water mirror of the reservoir. For the water renewal ratio and stream speed computations, the hydrodynamic simulations utilized the Environmental Hydrodynamics Base System (SisBAHIA®, http://www.sisbahia.coppe.ufrj.br/), a model used for natural water bodies under different scenarios [27]. The two-dimensional module of the FIST3D (Filtered in Space and Time 3D) hydrodynamic model was used to simulate water flow velocity and rate of water renewal during dry and rainy seasons. The FIST optimized spatial discretization allows for an optimal presentation of jagged contours and complex bathymetries. Bathymetric data was interpolated in the Surface Mapping System software (Surfer) using the Kriging regression method, with a spacing of 100 meters to obtain the Digital Terrain Model (DTM) for hydrodynamic modeling (S1 Fig). For the reservoir contour and spatial delimitation, the spatial discretization of the model was performed via biquadratic quadrangular finite elements. The water velocity was measured for both dry (October 23, 2020 to January 21, 2021) and rainy (April 1 to June 30, 2017) seasons for the entire area of the reservoir. The mesh has a grid resolution ranging from around 300 meters in the tributary rivers to approximately 2 km in the Três Irmãos Reservoir main body (S1 Fig).

Daily inflows from the NA reservoir and the arms where the Mato Grosso brook and Baixote stream are, and daily outflows of the TI dam and Pereira Barreto channel, in addition to the fluvial inflow of the tributary and the winds, were used as variables or constraints in the model. Due to the reservoir’s large size (∼132 km in length), a compartmentalized analysis of the rate of water renewal during a simulated three-month period was applied. The region was divided into 14 compartments (Fig 1B). The conservation of mass principle was applied to a generic substance diluted in the water mass with concentration C.

### Pilot sampling

Pilot sampling was conducted in September 2020 at one site in the reservoir (Fig 1A). To evaluate the best depth for sampling, three water samples were collected at the surface, middle, and bottom of the water column (1-, 13-, and 25-meter depths, referred to as P01, P13 and P25, respectively) (S1 Table). Three independent samples (R1, R2 and R3) were collected at each depth, resulting in nine samples. For each sample, one liter of water was collected utilizing a Van Dorn bottle, then stored in sterilized polypropylene bottles (Nalgon, Cat. No. 2330) at 8° C until filtered using 0.22 µm Sterivex filters (Merck-Millipore, Cat. no. SVGP01050) in 60 mL increments using disposable syringes. After filtration, the remaining liquid was expelled by injecting air, and the filters were stored at 4° C in sterile Longmire’s buffer (Tris 100 mM, EDTA 100 mM, NaCl 10 mM, SDS 0.5%) until the DNA extraction. Non-disposable equipment was cleaned with sodium hypochlorite solution (10% v/v) and rinsed with autoclaved distilled water to avoid cross-sampling contamination.

### Sampling stations in Três Irmãos Reservoir

After analyzing the data from both pilot sampling and the hydrodynamic modeling (see results section), 14 new stations were sampled in October 2021 during the dry season to assess the local biodiversity, hereafter called stations C1 – C4 and C6 – C15 (Fig 1A, S1 Table). Sampling stations covered each compartment used to evaluate water renovation ratios across the river. Stations C1, C2, C4, C6, C7, C9, C12, C13, and C14 were located in the main water body. Four stations were placed in areas that comprise the arms with (C3, C8, C10) and without tributaries (C11). One station (C15) was located at the entrance to the Pereira Barreto channel that connects the TI reservoir to the neighboring IS reservoir (Fig 1A).

Three replicates (R1, R2 and R3) were collected within each station, all 1 meter above the benthic bottom, resulting in 42 samples at different depths across among the 14 stations (S1 Table). The replicates were collected within approximately 10 minute intervals. All collection, processing, and storage procedures were the same as those described above for the pilot sampling.

### DNA extraction and sequencing

DNA extraction from the Sterivex filters was performed using the DNeasy^®^ Blood & Tissue kit (Qiagen) with adaptations [28]. The DNA concentration was determined using the Qubit dsDNA HS Assay Kit (Thermo Fisher Scientific). The amplicons from mitochondrial genes targeting the minibarcodes of cytochrome c oxidase I (COI), 16SrRNA (16S), and 12SrRNA (12S) were amplified and sequenced using the pair of primers described in S2 Table. Amplification was performed in 25 μl using Platinum™ Multiplex PCR Master Mix (Applied Biosystems), 1 μM of each primer, and 5 μl of DNA. The thermocycling conditions included an initial denaturation step at 95° C for 2 minutes and then 25 denaturation cycles at 95°C for 30 seconds each, annealing at different temperatures for each primer pair (48°C for CO1; 57°C for 12S and 53°C for 16S) for 90 seconds, and extension at 72°C for 30 seconds, followed by a final extension at 72°C for 10 minutes and finishing at 4°C. For the library preparation, the amplicons were indexed using Nextera XT indexes (IDT for Illumina). Library quality control was performed using Tapestation (Agilent). DNA concentration was normalized, and all amplicons were pooled in one library. The library was sequenced on Illumina NovaSeq 6000 S4 Flow Cell 2x150.

### Bioinformatic analysis

After sequencing, raw sequence reads were evaluated using FASTQC (http://www.bioinformatics.babraham.ac.uk/projects/fastqc) and MultiQC (https://multiqc.info/). All generated fastq files were imported into the QIIME2 pipeline [29] for further quality filtering and taxonomic classification. Primer trimming, read denoising, merging the remaining paired reads into amplicons, and chimera removal steps were carried out with DADA2 [30]. Here, all forward and reverse reads with less than 145 nucleotides overlapping were removed, as well as all reads with quality < 30 in Phred scale. The resulting Amplicon Sequence Variants (ASVs) were clustered into MOTUs (97% of identity among sequences) with DADA2. The taxonomy identification was performed using two different methods: 1) Naive Bayesian classifier [31] trained with the sequences’ dataset of the 12S Mitohelper [32], COI classifier version 4.0 [33] and the MIDORI 16S dataset [34]; and 2) BLASTn searches [35] using the curated mitochondrial reference database available in NCBI (https://ftp.ncbi.nlm.nih.gov/refseq/release/mitochondrion/), and the nucleotide non-redundant (nr) database [36]. Only hits with an e-value lower than 1e^-50^ and a query cover higher than 90% were allowed. ASVs and MOTUs that matched off-target species were discarded. Off-target species included non-eukaryotic organisms, human, domesticated animals (*Bos taurus*, *Canis familiaris*, *Felis catus*, *F. silvestris*, *Gallus gallus*, *Meleagris gallopavo*, *Mus musculus*, *Ovis vignei*) and marine animals (*Chimaera sp.*, *Coryphaena equiselis*, *C. hippurus*, *Scomber sp.*, *Homarus americanus*, *Rhinesomus triqueter*).

### Ichthyofauna communities

For the pilot sampling, a Binomial Generalized Linear Model (GLM) was employed to estimate the probability of encountering fish species at each depth level, allowing for the assessment of the optimal sampling design. Heatmaps of the log10 frequencies of MOTUs were constructed based on MOTUs for fish species. MOTUs frequencies were computed considering all three markers (CO1, 16S, and 12S) combined.

Based on fish MOTUs identified only up to the species level, ecological analyses of alpha- and beta-diversity were conducted in R. Two indices of alpha diversity were calculated with the vegan R [38] package within each locality: the Shannon Index (H’)[37], and richness (number of MOTUs). To compare the diversity indices across the 14 locations (beta-diversity), we constructed bar plot charts to visualize the species diversity in terms of fish orders. All graphs were constructed in R scripts using the *ggplot2* package [39].

### Evaluation of the eDNA-based biodiversity inventory completeness

To evaluate the completeness of our data, eDNA results were compared to the biodiversity data of TI built from Global Biodiversity Information Facility data (GBIF, https://www.gbif.org/) and a non-systematic literature review. In addition, both PCA and DAPC multivariate methods and rarefaction curve computation were employed. Using this integrative approach, we were able to assess the most informative samples and identify potentially redundant information. Only fish MOTUs identified up to the species level were considered, avoiding potential bias in biodiversity assessments.

The PCA was conducted to capture the variation among samples within the first two principal components (PCs), considering both MOTUs abundance and richness per locality. Individual and scree plots were generated using the factoextra package [39] from R v. 3.3.2 [41], respectively. The DAPC was implemented to cluster samples based on compositional similarity. Because DAPC maximizes differences among groups, it enables the exploration of sample similarities and the identification of potentially redundant samples in the experimental design. The optimal number of PCs was determined by the *optim.a.score* function from the *adegenet* R package [42], and genetic clusters were identified using the *find.cluster* function. Individual cluster assignments from DAPC were visualized using the *assignplot* function from *adegenet*. The contribution of each sample to the overall variability was assessed through computed eigenvalues within each PC.

The impact of reducing sampling effort on capturing local variation was tested through the creation of four sub-datasets, three of them lacking one of the three most contributing samples and one containing only the three most contributing samples. Rarefaction curves were then computed using all complete and sub-datasets using the function *specaccum* from the vegan R package [43].

## Results

### Hydrodynamic modeling shows a lentic environment and minimal water exchange in the TI Reservoir

Interpolation of bathymetry from the Três Irmãos (TI) Reservoir showed that the average depth of the region was 18.3 meters. The TI Reservoir is a predominantly lentic environment in both dry and rainy seasons, with more intense currents upstream, next to the Nova Avanhadava (NA) dam, and downstream near the TI dam. Hydrodynamic modeling results show that the water flows from the NA to the TI dam. The water velocity is higher near the main inflow into the reservoir at the NA dam (upstream in the reservoir) in both rainy and dry seasons (S2 Fig). Upstream, currents are more intense due to the topography of the channel, reaching up to 1.33 m/s and 0.56 m/s in the rainy and dry season, respectively. In the central region, the flow is very slow, with very low velocities (< 0.05 m/s) in rainy and dry seasons during all the modeled periods in the sections closest to the TI dam (S2 Fig). In the regions just upstream from the TI dam, currents are more intense than in the central region, but approximately 10% weaker than in the upstream region, presenting averages of 0.04 m/s in the rainy season and 0.02 m/s in the dry season.

Water renewal in the reservoir compartments varies between rainy and dry seasons. During floods, water renews faster, peaking at 50 days and then increases exponentially. Compartment 1 (closest to NA dam) renews 50% of its water in less than five days during the rainy season but takes 10 days in dry periods. In compartments 2, 4, and 6, a similar trend is observed for the first 50 days. After 70 days, sections 1, 2, and 4 have more than 95% of their water renewed during floods. Sections 1 and 2 in dry periods reach over 95% renewal in 90 days, while compartment 4 reaches about 80%. The furthest compartments (12, 13, and 14) only receive "new" water during the rainy season, renewing up to 10% after 50 days (S3 Fig).

The results indicated that the water flow in the arms tends to stagnate, regardless of the time of year, meaning there is a low exchange of water between the main flow and the arms. Taken together, the reservoir depth, water velocity, and water renewal rate supported the distribution of stations for water sampling, where they are allocated to capture eDNA in the different hydrological profiles of the reservoir and the arms.

### Bottom sampling and three-marker analyses improve the chances of fish eDNA capture

The pilot sampling obtained an average of 1.5 million of 150 bp paired-end reads per depth. After the read cleaning, amplicon merging, chimera removal, and clustering of identical amplicons, 615, 2,993, and 4,404 ASVs remained for 12S, 16S, and COI datasets, respectively (S3 Table). After applying the Bayesian classifier from QIIME2 and removing all off-target species, COI exclusively identified 24 taxa, while 16S found 3 and 12s, none. These datasets merged resulted in 27 non-fish taxa, which belonged to Rotifera, Copepoda, Annelida, Algae and Viridiplantae clades (S4 Table).

Regarding ichthyofauna, the GLM revealed that sampling at the benthic bottom (25 m) increases the chance of capturing fish eDNA (S4 Fig). COI, 12S and 16S detected five, four and one taxa, respectively, resulting in 10 taxa. Four species were identified by both 16S and COI, one by 12S and COI, and one by 12S and 16S. Only 4 species were detected by all three mini-barcodes (S4 Table).

### eDNA metabarcoding uncovers 171 MOTUs in the TI Reservoir, including 29 previously unrecorded species

As described in the Methods section, following the pilot assessments, a sampling campaign covering the entire reservoir through 14 stations was performed. After sequencing the 42 samples from 14 stations, 67.9 million paired reads for 12S, 31.1 million for 16S, and 74.9 million for COI were obtained (S3 Table). Following trimming and clustering steps, 2.9 million amplicons were identified for 12S, 22 million for 16S, and 14.8 million for COI. A total of 1187, 3687, and 7437 ASVs remained for 12S, 16S, and COI datasets, respectively. Subsequently, the classification using the Bayesian classifier from QIIME2 and removal of all off-target species, the ASVs clusters that share 97% of identity presented 34, 41, and 190 MOTUs, respectively.

The biodiversity inventory of TI Reservoir, combining historical and eDNA-based data, encompasses 577 taxa (S5 and S6 Tables). The dataset was divided into non-fish (S5 Table) and fish (S6 Table) to permit a broader view of the fishes detected, the main group of interest. The final dataset of non-fish metazoan resulted in 422 taxa, where 36 (∼8%) were also detected in this study (S5 Table). Of these, 20 taxa (∼5% of the 422) were exclusively detected using eDNA, with species belonging to Annelida (Clitellata and Polychaeta), Arthropoda (Copepoda, Ostracoda, Insecta, and Branchiopoda), Chordata (Avians) and Rotifera. The remaining taxa, previously reported in the literature, include Annelida, Arthropoda, Chordata, and Mollusca species (Fig 2 and S5 Table).

**Fig 2.**
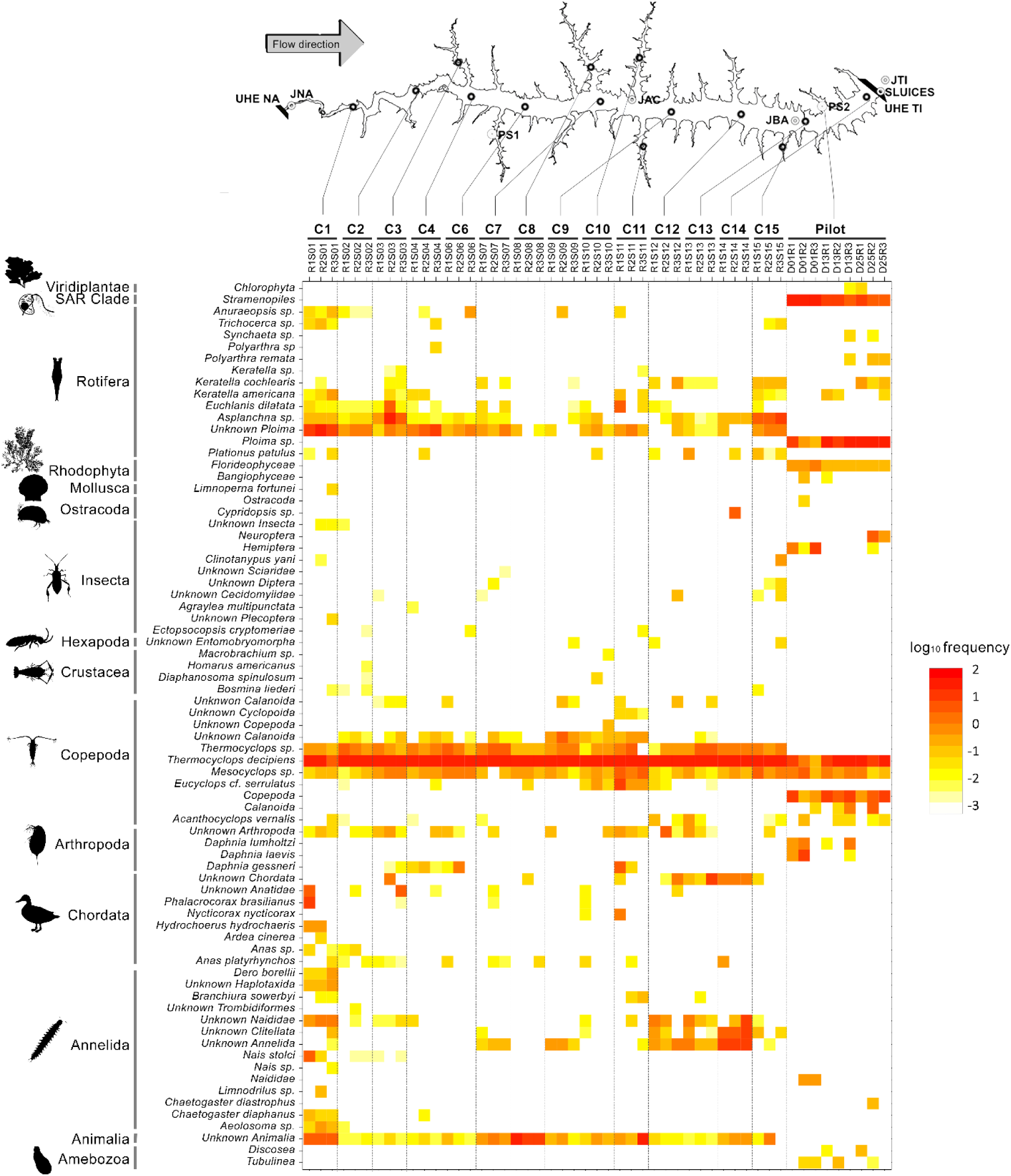
Non-fish taxa distribution and frequency across the Três Irmãos (TI) reservoir. Heatmap illustrating the distribution of non-fish Molecular Operational Taxonomic Units (MOTUs) identified by eDNA across sampling stations in the Três Irmãos (TI) reservoir. The TI reservoir is shown at the top with all sampled stations C1-C4 and C6-C15. Stations abbreviations as in S1 Table. Hydroelectric power plants Três Irmãos (TI) and Nova Avanhandava (NA) are shown at the extremes of the reservoir. The yellow to red color spectrum represents the log10 transformation of the average frequency. Solid black lines on the left delineate the observed phyla.

Arthropods, especially copepods, were highly abundant across all stations, including the pilot station. We also found four non-fish taxa never previously reported in the TI reservoir, even though they had been reported in upstream reservoirs and other rivers of the Paraná Basin: an Olygochaeta (*Chaetogaster diastrophus)*, a copepod (*Acanthocyclops vernalis)* and two rotifers (*Polyarthra remata* and *Synchaeta* sp.). SAR (Stramenopila, Alveolata, and Rhizaria) MOTUs belonging to the Stramenopila clade were not identified beyond this level because of the lack of reference sequences.

Metabarcoding eDNA matched 12 species previously identified in TI through traditional taxonomy: two annelids (*Branchiura sowerbyi* and *Limnodrilus* sp.), one arthropoda (*Diaphanosoma spinulosum*), three birds (*Nycticorax nycticorax*, *Mycteria americana* and *Phalacrocorax brasilianus*), one mammal (*Hydrochoerus hydrochaeris*), one mollusc (*Limnoperna fortunei*), and four rotifers (*Asplanchna* sp., *Euchlanis dilatata*, *Polyarthra* sp. and *Trichocerca* sp.). Although some taxa remained unclassified, most belong to groups previously found in TI and other rivers of the Paraná Basin (S6 Table). Nonetheless, a few species present in previous reports in the literature were not detected through DNA sequences obtained in this work, such as Cyanophyta, predominant during 2011 [43]. No macro-crustacean was detected, including the exotic *Macrobrachium amazonicum* and *Macrobrachium jelskii*, both previously found in TI [45]. Insects [46], Molluscs [46], except for *L. fortunei*, and Platyhelminthes already reported in this area were not detected either.

For the ichthyofauna, the resulting inventory comprised 135 species (S6 Table), with 35 species being identified through both literature review and eDNA surveys. In comparison, 81 species (∼60%) were only documented in the literature over the past 50 years. Nine species detected with eDNA metabarcoding had not previously been recorded in TI: one cypriniform (*Gobio gobio*), seven characiforms (*Hemiodus orthonops*, *Hoplias aimara*, *Hyphessobrycon erythrostigma*, *Leporinus piau*, *Schizodon fasciatus, Schizodon knerii, Metynnis melanogrammus*), and one cichliform (*Crenicichla cyanonotus*), here found in multiple sites (Fig 3). Approximately 60% of all species were captured through both eDNA and traditional monitoring in 2020-2021, including *Roeboides descalvadensis*, *Hoplias malabaricus*, *Crenicichla semifasciata*, *Cichla kelberi, C. piquiti, Satanoperca pappaterra, Serrasalmus maculatus, Oreochromis niloticus, O. aureus* and *Plagioscion squamosissimus*, for instance. Only 19 species (∼14%) had GBIF records for the TI reservoir.

**Fig 3.**
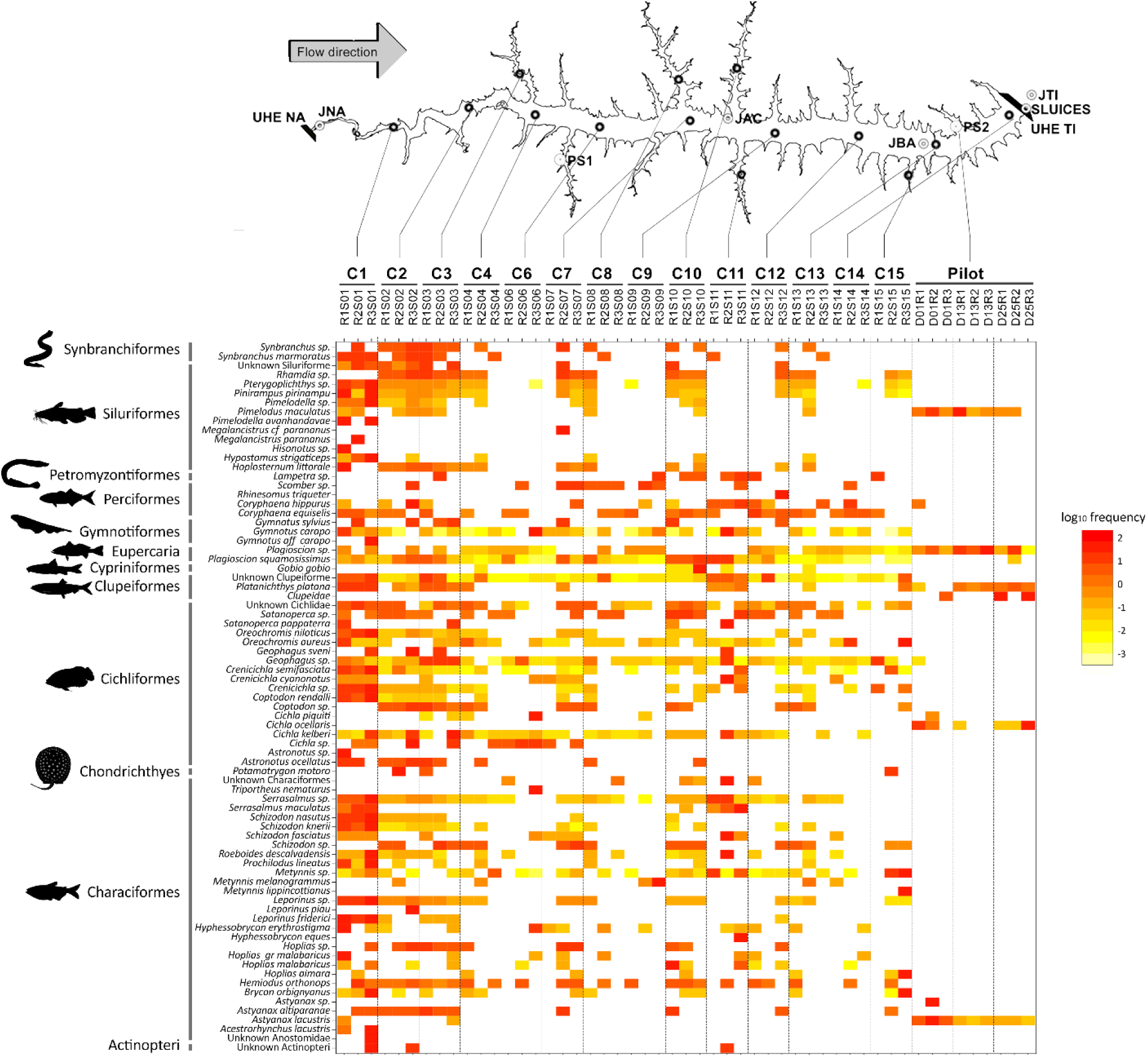
Ichthyofauna distribution and frequency across the Três Irmãos (TI) reservoir. Heatmap illustrating the distribution of fish Molecular Operational Taxonomic Units (MOTUs) identified by eDNA across sampling stations in the Três Irmãos (TI) reservoir. The TI reservoir is shown at the top with all sampled stations C1-C4 and C6-C15. Stations abbreviations as in S1 Table. Hydroelectric power plants Três Irmãos (TI) and Nova Avanhandava (NA) are shown at the beginning and at the end of the reservoir. The yellow to red color spectrum represents the log10 transformation of the average frequency. Solid black lines on the left delineate the observed orders.

Seventy-seven fish species were identified through eDNA when considering all locations from both sampling and station samples (Fig 3). These species are distributed among nine orders, namely Cichliformes, Characiformes, Chondrichthyes, Cypriniformes, Eupercaria, Gymnotiformes, Myliobatiformes, Siluriformes, and Synbranchiformes. The Cichliformes and Characiformes orders were the most abundant in almost all stations (Fig 4A), followed by Gymnotiformes, represented here by species of the genus *Gymnotus*, exhibiting significant presence, especially in station C01. The order Clupeiformes, represented by a single species (*Platanichthys platana*), was found in the pilot sampling, and identified in three other stations (C03, C04, and C15). Chondrichthyes species were detected in several stations, but in low frequencies, ranging between 3% and 8% per sample.

**Fig 4.**
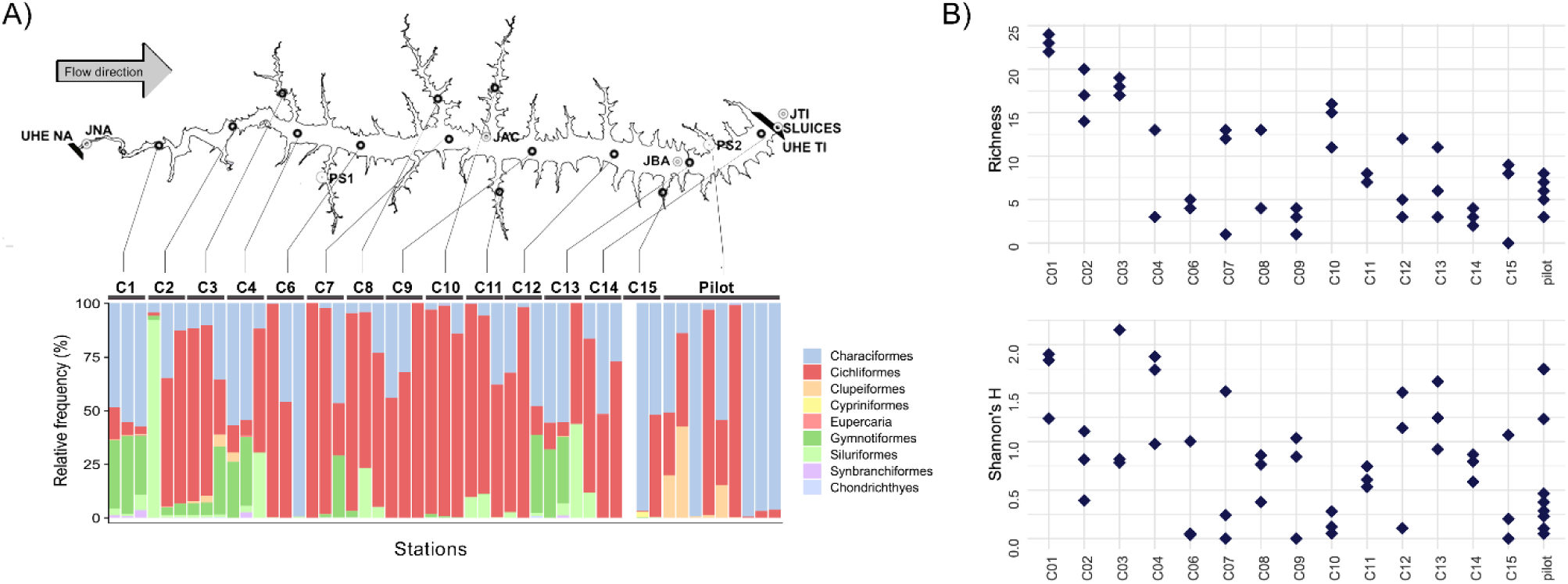
Alpha diversity results for ichthyofauna taxa detected in the Três Irmãos (TI) reservoir. Based on fish Molecular Operational Taxonomic Units (MOTUs) identified through eDNA per sampling station, A) the barplot displays all orders found within the Três Irmãos (TI) reservoir. Colors indicate the orders as shown in the legend. The TI reservoir is shown at the top with all sampled stations C1-C4 and C6-C15. Stations abbreviations as in S1 Table. B) Richness and Shannon’s H index (y-axis) were computed for each station (x-axis). Each diamond represents one sample collected at the respective station.

Station C01 exhibited higher variability in terms of the number of taxa and also had higher MOTU frequencies compared to all other stations, revealing some exclusive species, such as *Acestrorhynchus lacustris*, *Pimelodella avanhandavae*, and *Gymnotus carapo*. The only autochthonous species identified was the Siluriforme *Pimelodus maculatus*. The other six species are allochthonous (originally from other basins) such as the *Plagioscion squamosissimus* (Corvina) and the three cichliforms (*Cichla piquiti*, *C. kelberi*, and *C. ocellaris*) all native to the Amazon.

From the six native fish species known to be introduced annually into the TI reservoir as part of a local restocking program, *Brycon orbygnianus* (Piracanjuba) and *Prochilodus lineatus* (Curimbatá) were detected by eDNA metabarcoding at nine and seven sampling stations, respectively. The other four species that are part of the restocking program *Piaractus mesopotamicus* (Pacu-guaçu), *Pseudoplatystoma corruscans* (Pintado), *Salminus brasiliensis* (Dourado) and *Leporinus elongatus* (Piapara) were not detected with eDNA.

### Diversity variation among samples from the same station is greater than across stations

Despite being collected only minutes apart, diversity indexes, i.e. richness and Shannon’s H, of the ichthyofauna communities showed greater variation among samples from the same station than across stations (Fig 4B). For instance, the C12 station presented richness in one of the replicates that was almost two times greater than the other two replicates (12 compared to five and three). Stations C01, C02, and C03 exhibited the greatest richness, whereas the highest Shannon’s H values were observed in stations C03, C01, and C04, respectively.

### A quarter of the sampling effort was sufficient to reach the biodiversity plateau of the TI reservoir

A PCA was built with only the fish MOTUs identified to the species level. The individual position in the graph indicated a high similarity in composition among the samples, except for those from station C01 (C01R1, C01R2, and C01R3) and a few from stations C02 (C02R3), C03 (C03R1) and C07 (C07R2) (Fig 5A). These samples were collected at stations considerably distant from all others, potentially suggesting the presence of species exclusive to these sites.

**Fig 5.**
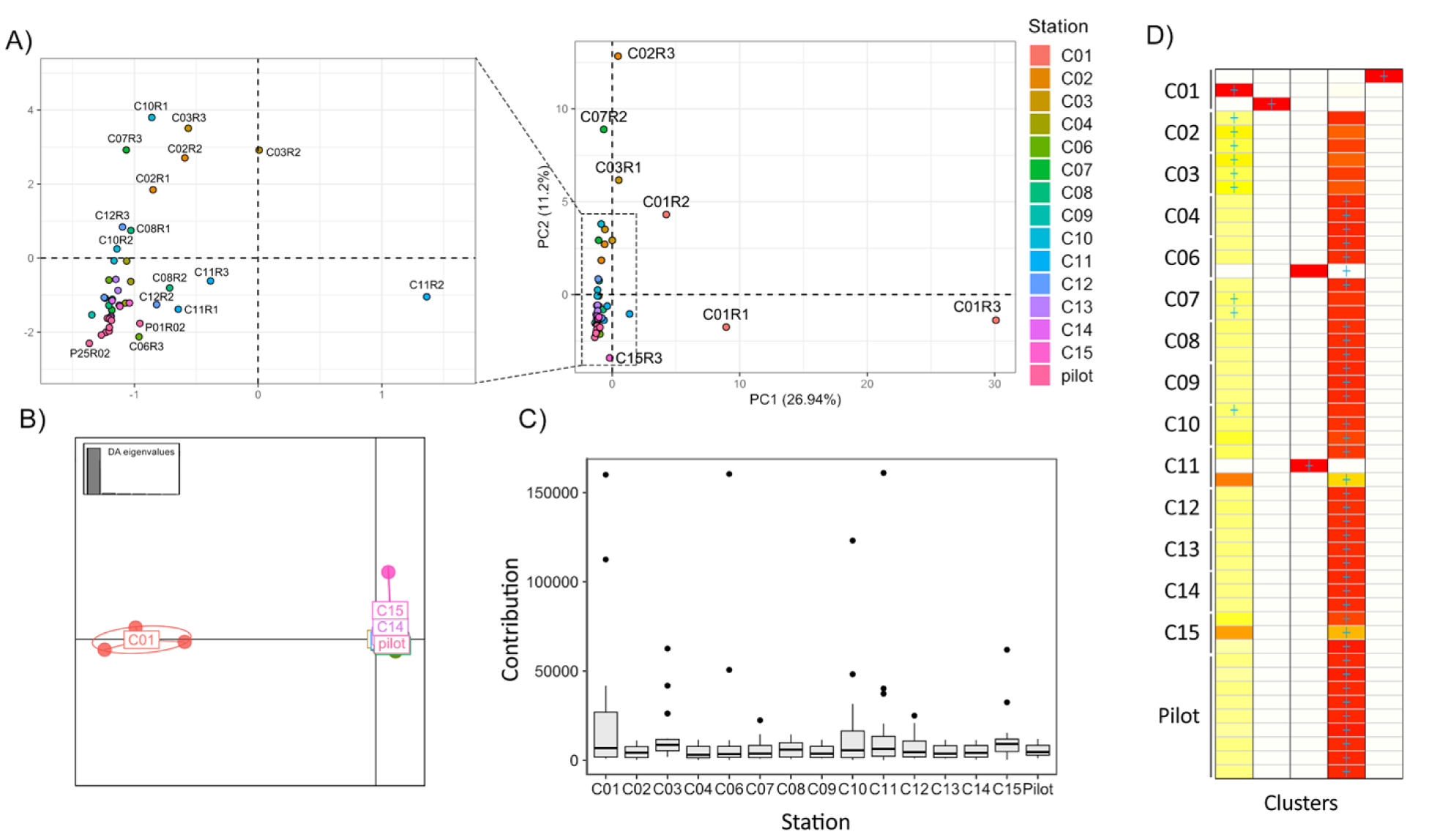
Beta diversity results for ichthyofauna taxa detected in the Três Irmãos (TI) reservoir. Based on fish Molecular Operational Taxonomic Units (MOTUs) identified through eDNA per sampling station, A) Principal Component Analysis (PCA) illustrates the variance in community composition. Each dot represents a sample from a station, with colors designating stations, as in the legend. Stations abbreviations as in S1 Table. Samples plotted close to each other are zoomed in the minor graph on the left side. B) Discriminant Analysis of Principal Components (DAPC) exhibits the maximization of differences among groups (6 PCs, K = 5 according to the BIC method). The colors correspond to those in PCA. C) Boxplot shows the contribution of each station and their samples to the segregation into five clusters. D) Assign plot illustrates each individual sample’s assignment, considering K = 5. Darker shades of red indicate a higher chance of the sample belonging to that cluster.

In contrast, no distinct clustering pattern was observed for all other sites, indicating that despite the pilot sampling being conducted 12 months earlier, the composition remained very similar, preventing the segregation of these samples in the ordination space. However, the maximization of differences between these groups reinforces the pattern observed in the PCA, where only C01 appears different from all others, and the segregation among the remaining locations becomes indistinguishable in the ordination space (Fig 5B). When analyzing the sums of contributions from each station to the six principal components, stations C01, C10 and C11 exhibited the highest indices (Figs 5C and S5). By using six PCs according to the alpha score results, which reached 98.7% of the total variation, the DAPC identified five distinct groups: three groups comprised of one sample each from station C01, one group comprised of one sample each from C06 and C11, and the fifth group comprised of all the remaining samples (Fig 5D). In addition, a single sample from station C15 presented equal chances of belonging to two distinct groups.

The composition graph of all five clusters revealed that samples from C01 belong to different clusters, suggesting that even being sampled within the same station, the three obtained samples were different enough to be considered distinct community groups (Fig 5D). After these findings, using the samples with the highest contribution for the six PCs, rarefaction curves were computed for the datasets: i) containing all samples; ii) without C01; iii) without C11; iv) without C10; and v) containing only C01, C11, and C10 samples. For the complete dataset, as well as for those lacking one sample, the curves reached the plateau between 10-12 samples. The graphs indicated minimal changes in the plateau area, with slightly more noticeable differences observed when removing C01, suggesting that additional samples would be needed to fully capture the observed diversity (Fig 6). When C10 and C11 were removed, the results were relatively similar and did not exhibit a clear alteration in the curve shape. The rarefaction curve built using only C01, C10, and C11 samples almost reached the plateau area after nine samples, the total number (S5 Fig).

**Fig 6.**
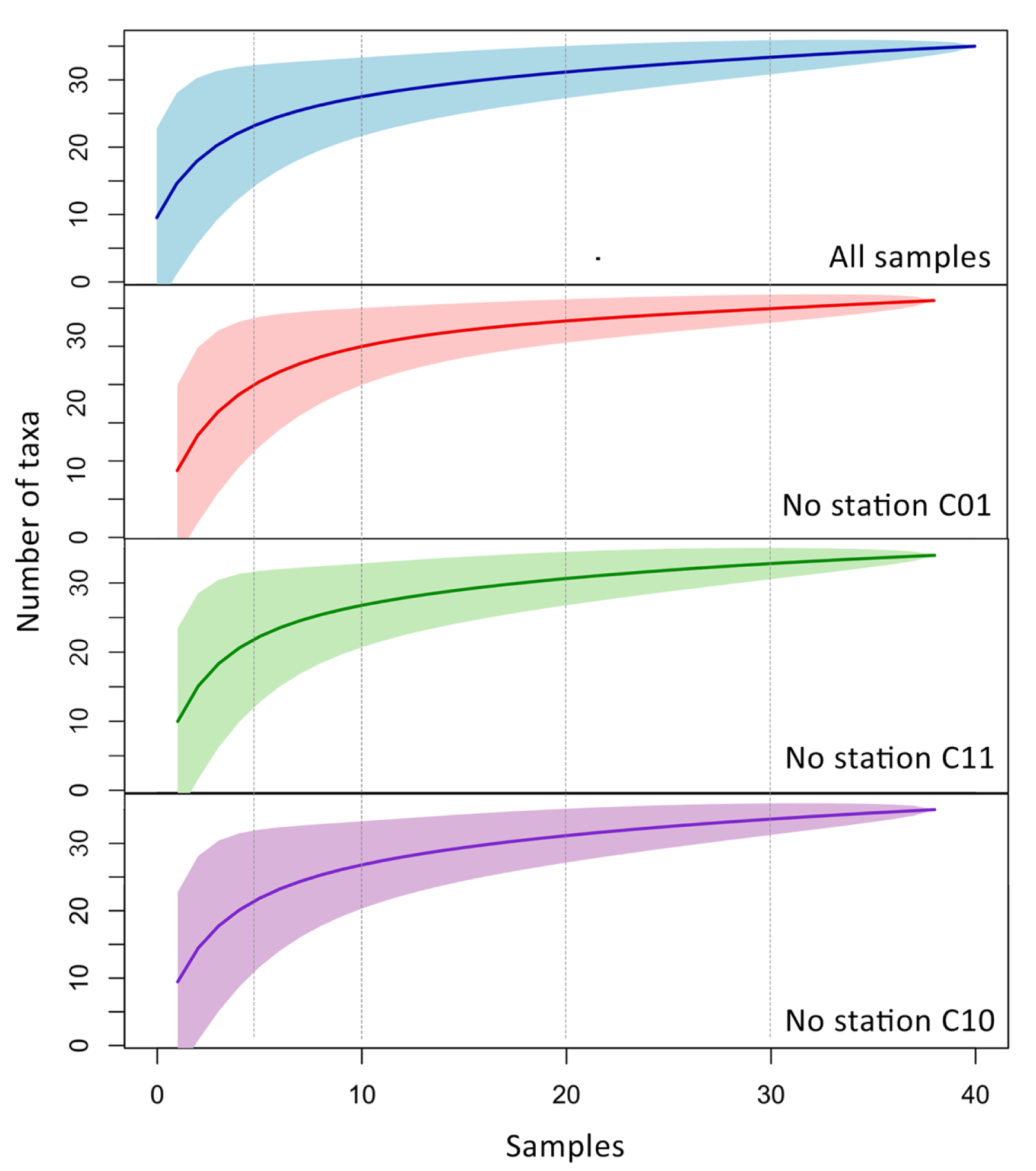
Rarefaction curves constructed based on fish richness using distinct sub-datasets. From top to bottom, the curves represented were built using all data, without station C01, without station C11 and without station C10, respectively. The number of taxa is indicated in the y-axis, and the number of samples (42 in total), in the x-axis.

## Discussion

This study provides an important insight into the optimal sampling strategy when conducting eDNA surveys in hydroelectric power plant reservoirs. Here, through the combination of hydrodynamic modeling and pilot sampling, we could delineate an optimal experimental design for the TI reservoir. In this study, we managed to perform a biodiversity baseline of a large reservoir for the first time in South America.

Metabarcoding successfully detected a substantial number of species previously documented in the literature over the past 50 years, along with nine fish species never reported in the reservoir before. We found that larger sampling sizes might not be the best approach when assessing biodiversity in lentic reservoirs using eDNA. Instead, it appears preferable to sample fewer locations with more replicates. Moreover, a twelve-month interval between sample collections did not seem to significantly influence diversity measures or the detection of species. Lastly, our data provides a valuable contribution to future biodiversity monitoring studies in reservoirs, aiming to achieve optimal utilization of resources with lower costs and better cost-effectiveness through molecular data.

This study contributed 20 new previously unreported non-fish species for the TI area (∼5%). Overall, there was a high diversity of non-fish taxa, with annelids and rotifers found across several samples (Fig 2). Copepods, on the other hand, were abundant but less diverse, mainly represented by the genera *Thermocyclops* and *Mesocyclops*. Chordates were predominantly represented by birds with marine habitats, such as herons (*Ardea cinerea*) and Anatidae species (ducks, geese, and swans), found just upstream from the dam (C01-C03) and in the middle of the reservoir (C04, C07, C10, and C12). The only non-avian chordate detected was the capybara (*Hydrochoerus hydrochaeris*) in C01, a mammal commonly found in riverside and lakeside areas [47]. Contrary to what would be expected, the invasive golden mussel species *Limnoperna fortunei*, highly abundant in the TI HPP intake [48], was detected only in C01. The animal’s body size and the primers chosen have direct influence on the eDNA survey; low DNA concentration [49] or the lack of amplification [16] or could explain its absence. Most non-fish species are microscopic or from meiofauna, with a body size smaller than 1 mm, such as rotifers, copepods, ostracods, and a few annelids. Therefore, these results reinforce how eDNA plays an important role when it comes to the identification of smaller bodies or reduced biomass animals, as already pointed out by previous studies [5].

Regarding the ichthyofauna, the eDNA survey identified nine species unreported to date, which corresponds to a ∼6% increase in all species in the inventory of the study area. Nine major fish orders were detected across all samples, reinforcing previous reports for the TI. We identified abundant carnivorous species such as *O. aureus*, *O. nilotocus*, *C. piquiti*, *C. kelberi*, *C. ocellaris* and *Plagioscion squamosissimus*. In fact, three of the most abundant species captured by traditional monitoring during 2021 [50] were also among the most frequently detected at multiple stations using eDNA (*P. squamosissimus* and *C. kelberi*), supporting our findings.

We also detected two native *Crenicichla* species, one of the richest groups in terms of species among the South American cichlids [51]. From the six taxa never previously reported in the TI reservoir, *Hemiodus orthonops* was detected at high frequencies at all sampling points. With regard to the distribution across compartments and communities’ patterns as characterized by eDNA, it is tempting to look for an ecological explanation for the presence or lack of some species in certain locations. However, several factors like species occupancy [52, 53], eDNA persistence and quantity [54, 55], primer bias [16] and DNA concentrations [49] may influence the distribution pattern of eDNA. Therefore, the absence of a species should not be solely attributed to its lack of occurrence. Following the same logic, the frequency of DNA fragments in the heatmap represents a rough estimation of the abundance of the species. For this reason, the eDNA abundance should be interpreted with caution as it may reflect body size of individual species [56] or drift [56], even for fishes [58].

When compared to the short-term traditional monitoring conducted during 2020-2021, eDNA successfully detected 60% of all fish taxa captured using traditional methods and more than 80% of all fish genera identified through traditional surveys (S6 Table). Several studies comparing eDNA metabarcoding and established monitoring methods have been conducted in rivers [19, 59, 60] or estuaries [51]. Consistent with our findings, most of these studies have shown that eDNA methods exhibit similar performance compared to capture-based fishing methods, but only for short and medium-term monitoring. For example, Goutte *et al.* (2020) found that eDNA was more efficient than traditional surveys lasting 3 to 6 years, but not in comparison to 14 years of monitoring efforts.

In traditional community ecology, which relies on specimen counting, the distribution of organisms is usually clustered under the influence of environmental and biological factors such as physical barriers, nutrient availability and competition. However, eDNA sampling is not bound by the same constraints. Instead, it is influenced by drift, sedimentation, and degradation rates [5,54, 56, 58] and, once released, eDNA tends to homogenize in the water flow, resulting in a gradient-like distribution that becomes more uniform over time. These processes effectively blur the boundaries of the originally distinct organismal clusters, illustrating the distinct dynamics between direct ecological observations and molecular ecological data. Here, eDNA could also detect species never reported for the study area, compensating for some of the limitations of the traditional monitoring methods, such as erroneous identification, dedicated time, and seasonal changes [61]. These findings support eDNA metabarcoding as an effective method when evaluating fish communities, as already affirmed in previous studies [62]. For this reason, the eDNA can be employed for rapid and large-scale biodiversity assessments for monitoring programs [17], reducing costs and minimizing interventions in aquatic environments.

The employment of multivariate analysis combined with rarefaction curves allowed us to assess whether the eDNA-based diversity was robust and to identify potential redundant sampling sites while highlighting the most diverse locations. Our findings show that even though the water samples were collected at the exact same location within a few minutes’ interval, the eDNA obtained was different enough to distinguish these water samples. Both alpha diversity analysis and PCA showed that many samples within the same stations are highly divergent, presenting distinct numbers of taxa and do not contain similar variance regarding community composition (Figs 4 and 5A). The PCA revealed that station C01 contributed the most to the observed variation (Fig 2A). This overrepresentation appears to result from species exclusive to this compartment, as supported by the frequency heatmaps (Fig 3). The C01 station alone accounted for 37 of 79 non-fish MOTUs (∼46%) and 32 (80%) of all fish species. However, the three samples from C01 presented higher variation than the expected from replicates, as observed in the DAPC clustering analysis. Due to potential differences in composition and abundance, two out of the three samples from C01 are distinct enough to be considered a separate ecological group (Fig 5D). For C01, a potential explanation for these results is the higher current speed (0.5 m/s compared to 0.05 m/s in other stations). Because eDNA behaves like fine particulate organic matter, the concentration of eDNA can be influenced by the hydrological profile, being transported according to the velocity of water movement (60). Consequently, the eDNA obtained during each sampling would substantially differ from each other. However, it is also possible that the ichthyofauna habitat preferences are shaping these communities’ differences. As previously demonstrated, the ichthyofauna exhibits distinct preferences regarding water velocity and depth, for example, in lotic systems [63, 64].

It is worth noting that in the PCA, all pilot sampling was very close to all stations except C01 (Fig 2A). This indicates that despite the twelve-month difference between the samples, this time gap did not result in significant differences in the composition of local communities. However, the different sample sizes from the two samplings, plus the short duration of the study, highlights the need for more comprehensive spatial analyses to confirm the time-scale effects on fish communities in the TI. Previous studies have indicated significant changes in community composition after a few weeks [65]. In some cases, temporal dynamics *per se* are not the most determinant factors; rather, environmental factors such as seasonality, salinity, or precipitation can have a greater influence on species distribution and occurrence [61].

Following station C01, stations C10 and C11 demonstrated significant contributions to the clustering analysis of groups from DAPC (Fig 2D). Interestingly, these samples are at the boundaries (C01), and at the center arm (C10 and C11) of the TI reservoir (Fig 1A). To assess the impact of sampling design with and without these stations, we constructed rarefaction curves to determine the number of samples to achieve our observed number of MOTUs. When any of the C01, C11, or C10 samples were excluded, the plateau area of the curve did not change significantly, remaining around 10 to 12 samples (Fig 6). However, when using only these three stations, the rarefaction curve almost reached the plateau shape in a smaller sample size (∼8 samples) (S5 Fig). These data show that our sampling effort with 42 samples and 14 stations was excessive, resulting in a considerable amount of redundant information. For this reason, we suggest that for the TI reservoir, a sampling size with only three strategically positioned stations with more samples within each one would be capable of capturing the same number of MOTUs as 42 samples within 14 stations. The location with higher diversity was the one with higher current speed (C01), supporting that the observed diversity does not follow a geographic pattern per se and, instead, it can be a result from either abiotic or biotic factor iterations, still unknown. Either way, the observed diversity could be expected at random sampling with at least 12 sites across the entire reservoir.

When considering each replicate as a sample, the diversity is high and multiple species could be detected. However, the species distribution is very similar among them. Therefore, increasing the sampling effort by adding replicates in fewer locations could increase the number of taxa. It is not the first time that increasing the number of replicates was suggested for river systems. For instance, Shaw *et al.* [19] already emphasized the same discussion when evaluating a river area in South Australia, where the authors recommend more replicates to increase the chance of detecting DNA fragments from taxa that are patchily distributed, with low abundance, low biomass or rare occurrence.

## Conclusions

We constructed a large-scale biodiversity inventory to evaluate the diversity of a reservoir’s communities, focusing specifically on metazoans and ichthyofauna. Through the use of various descriptive and statistical metrics, we assessed the robustness of our biodiversity inventory and determined the optimal experimental design. We found that conducting consistent sampling at fewer sites with more replicates was more effective in capturing biodiversity. Metabarcoding proved to be a valuable tool, corroborating previous reports based on traditional monitoring surveys and identifying several previously unreported species.

## Acknowledgments

We thank Angélica Beccato, Environment and Land Manager at Tijoá Energia, for her support and collaboration in developing this work.

## Data Availability

All sequences generated in this study are available on NCBI SRA (BioProject PRJNA1105563).

## Author Contribution

TC performed data curation, formal analyses, methodology implementation, and writing for this study. ALQT also conducted data curation and formal analyses, along with applying the methodology. MC and GZ contributed to the methodology approach and formal analysis of hydrodynamic modeling, as well as its visualization. MG and HBP provided formal analyses and validated the generated data. CMP, GMC, and LTAG contributed to the validation and discussion regarding the interpretation of the results. JAA, DLASA, and MFR were responsible for the conceptualization, funding acquisition, and supervision of this study. All authors reviewed and contributed to the writing of this manuscript.

## Supplementary information

**S1 Fig.**
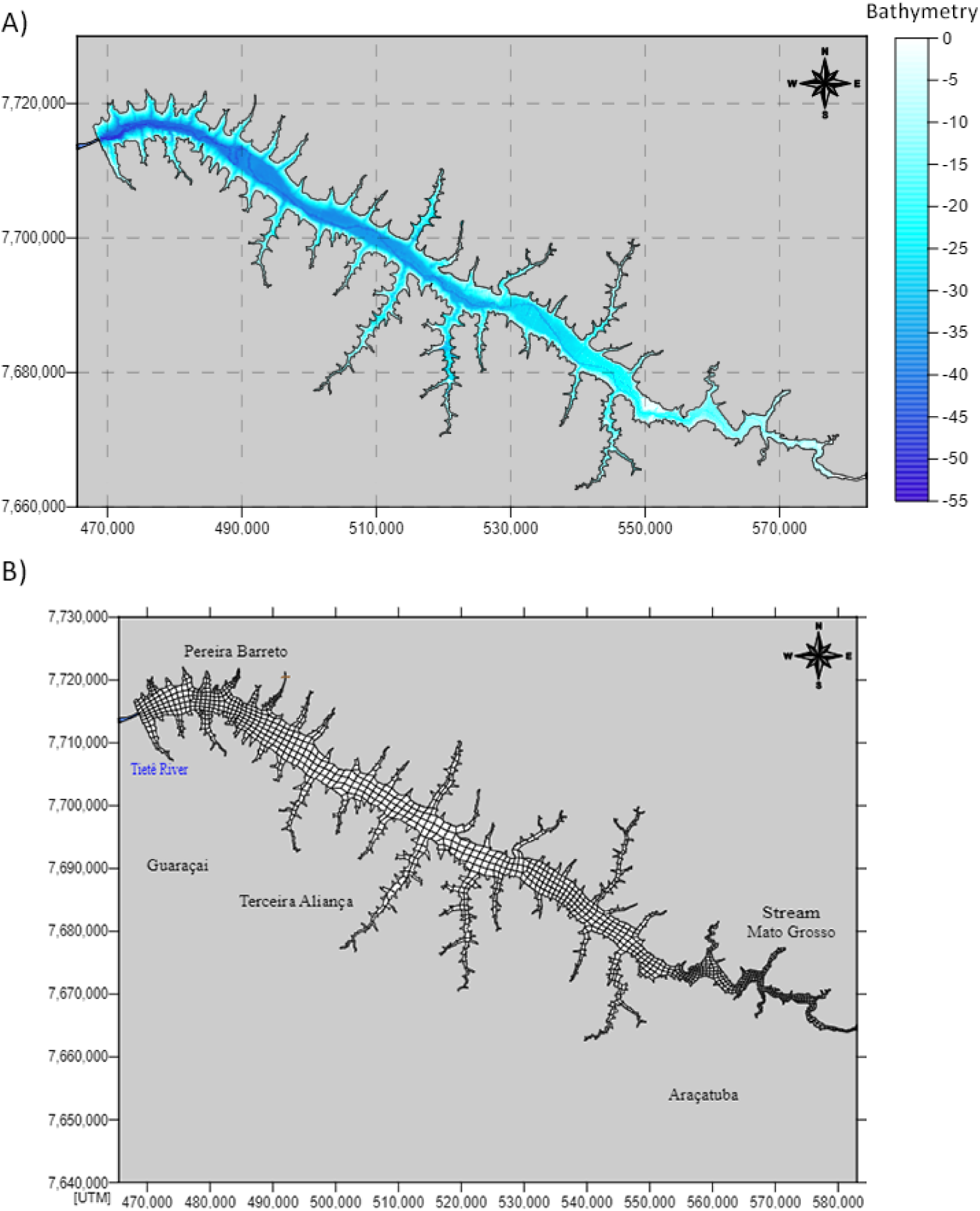
Metrics and spatial information regarding the Três Irmãos (TI) reservoir for hydrodynamic modeling. A) Bathymetry distribution across the reservoir. Darker shades show deeper sites. B) Biquadratic quadrangular finite elements for spatial discretization of the model.

**S2 Fig.**
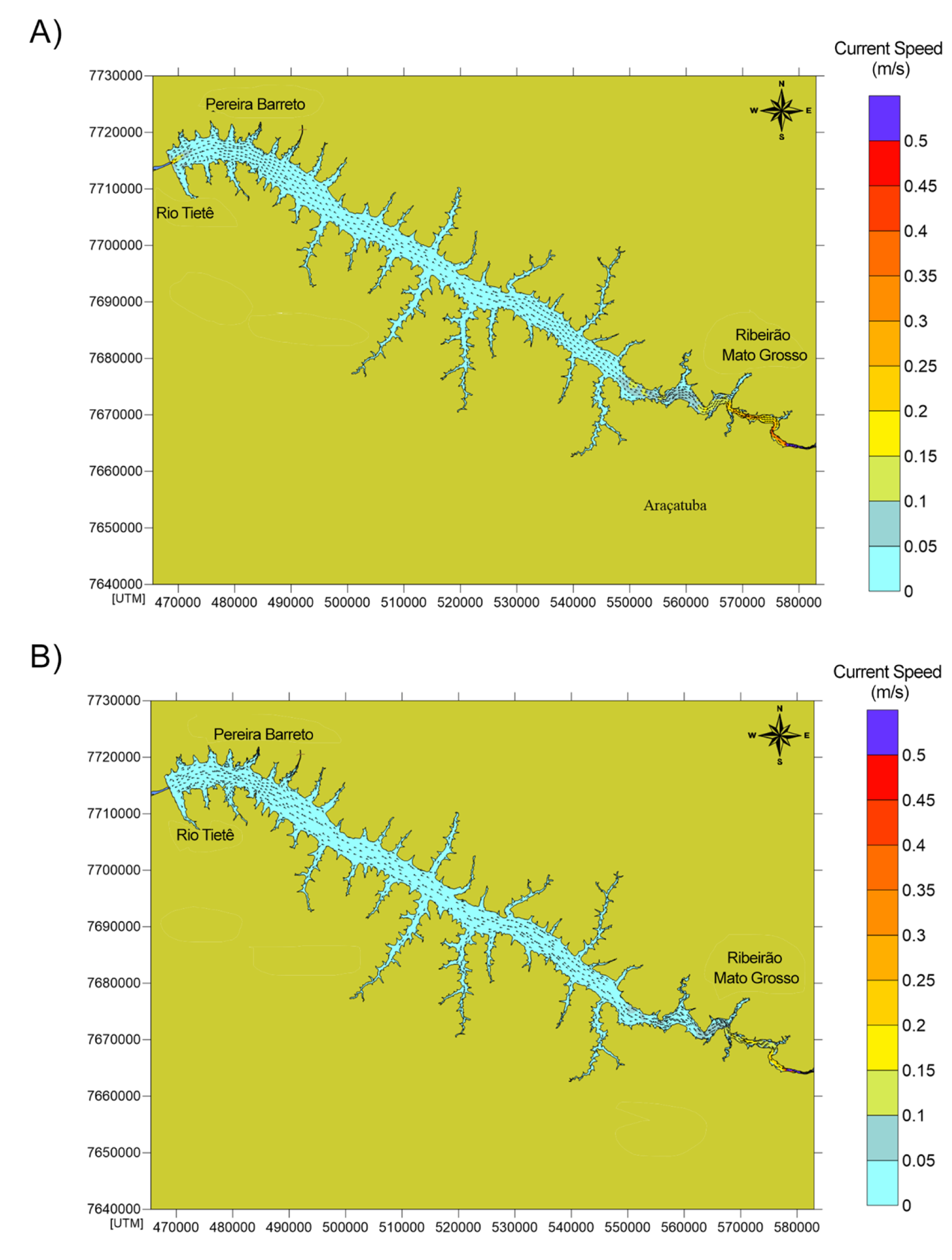
Hydrodynamic modeling for the entire Três Irmãos (TI) reservoir. Maximum current speed as measured during the A) rainy and B) dry seasons.

**S3 Fig.**
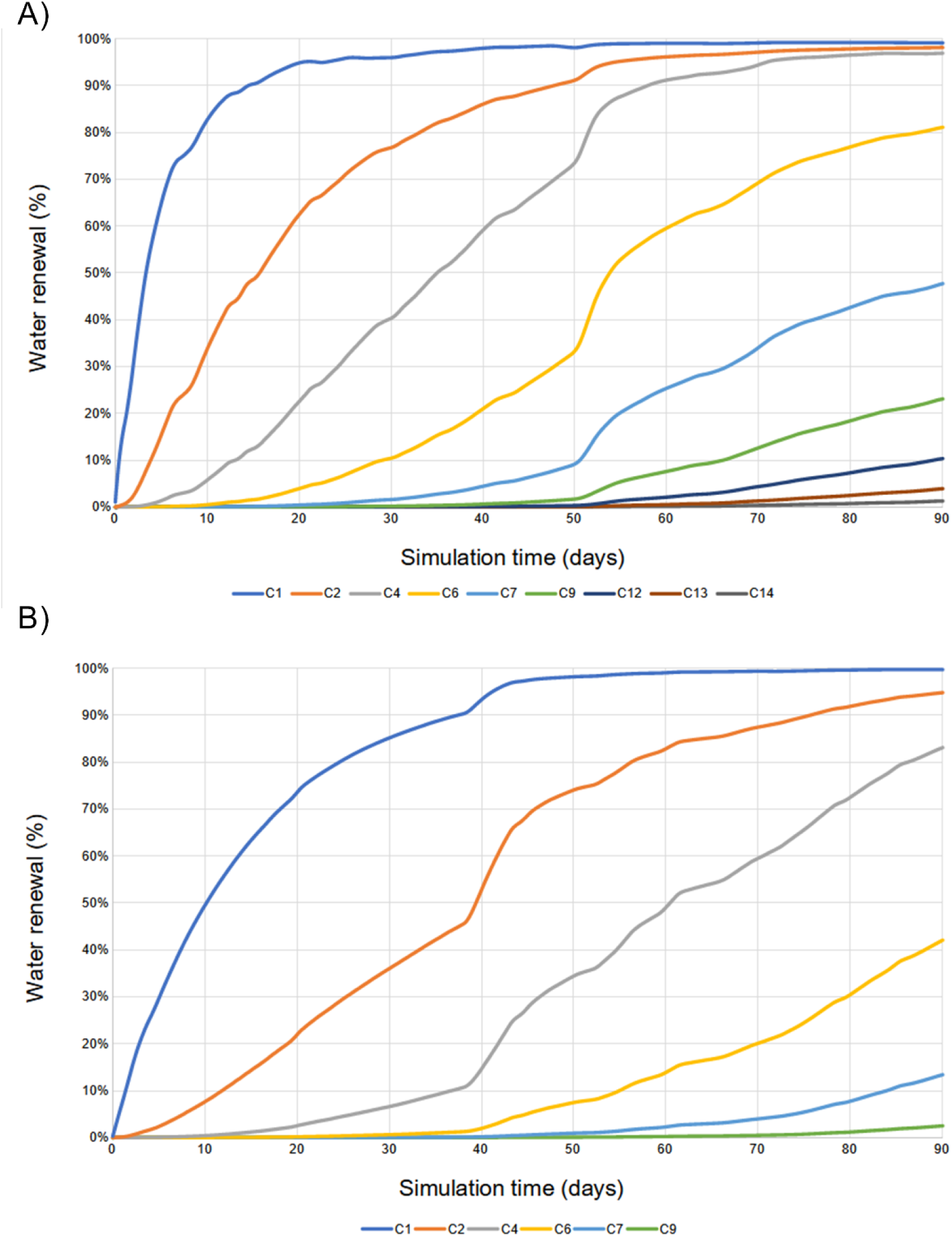
Computed water renewal rate for the Três Irmãos (TI) reservoir. The water renewal is shown moving from Nova Avanhandava to Três Irmãos reservoir during A) rainy and B) dry scenarios.

**S4 Fig.**
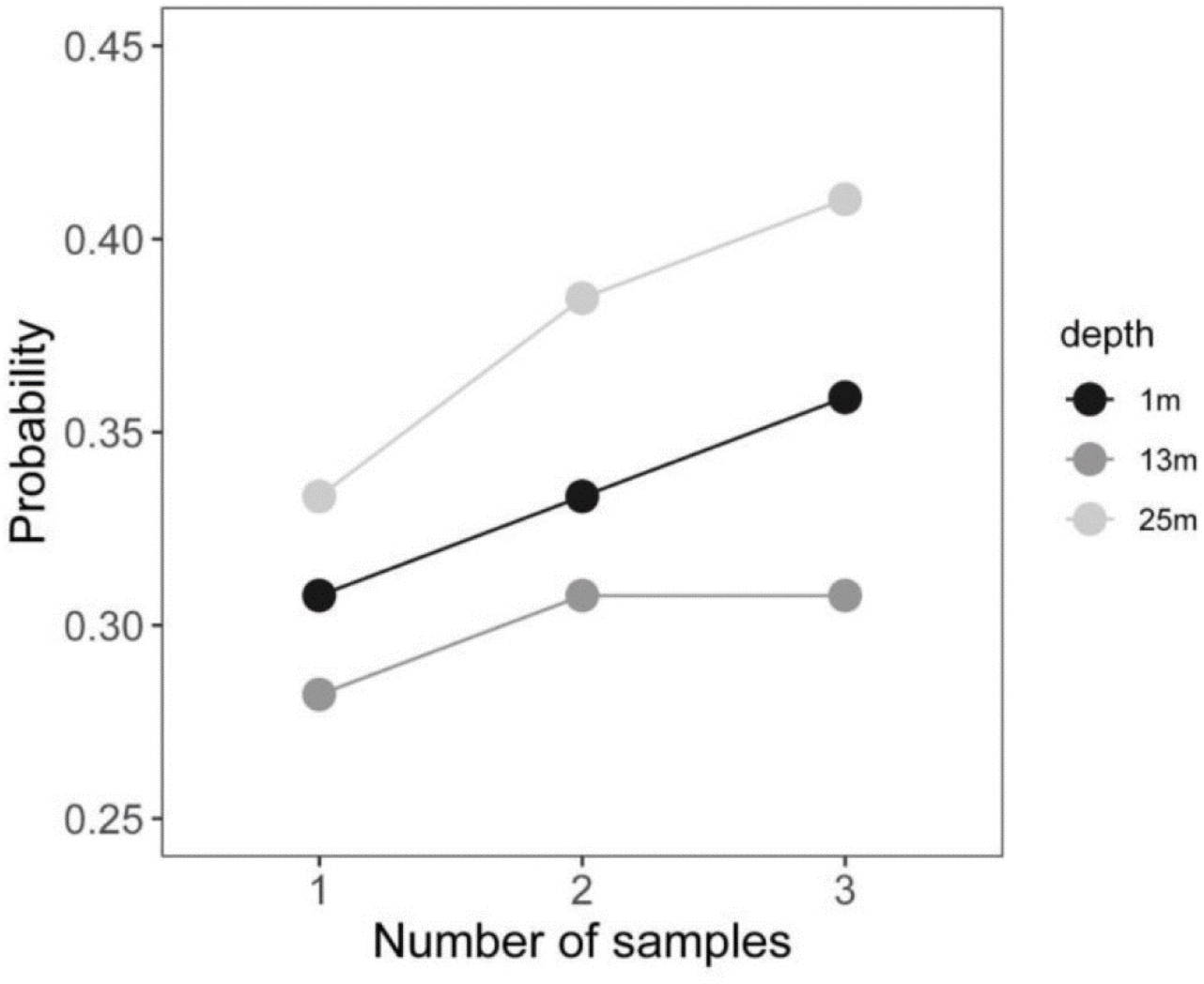
Probability of identifying fish by sampling for eDNA at different depths. Generalized Linear Model (GLM) showing the probability (y-axis) of sampling fish taxa in different depths (1 meter, 13 meters and 25 meters from the surface) after three samples (x-axis). Each line represents one depth according to the colors in the legend.

**S5 Fig.**
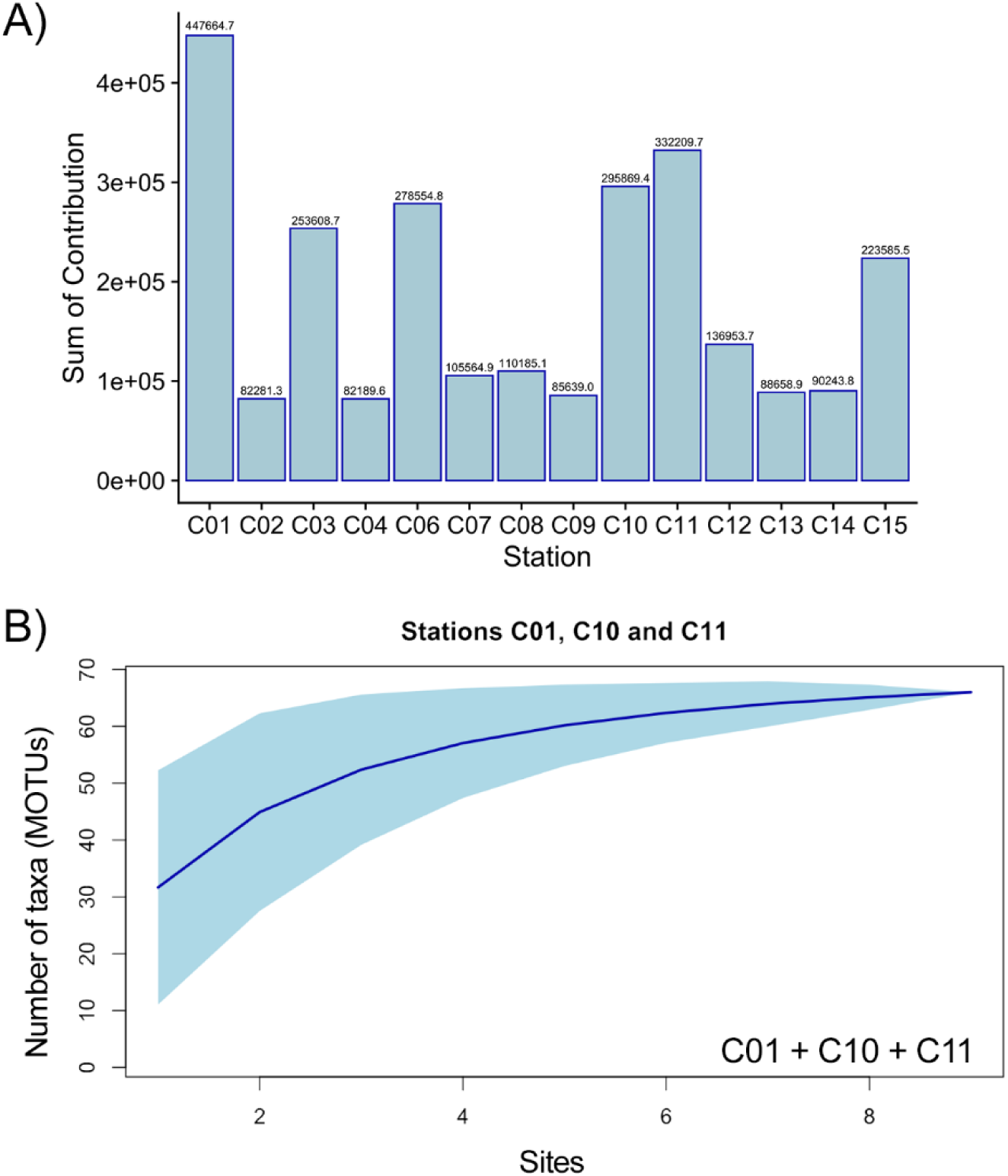
Statistics from the Discriminant Analysis of Principal Components (DAPC). A) Sum of contribution of each sampled station for the DAPC clustering assignment. Given the differences in sampling size, the pilot sampling is not included. Station abbreviations as in S1 Table. Each bar corresponds to one station, and the value shows the sum of contribution for the six first Principal Components (see methods). The stations with the highest values are C01, C11 and C10, respectively. B) Rarefaction curve constructed using only samples from C01, C11 and C10, where the curve reaches the plateau curve around eight samples.

**S1 Table. Sampling locations’ information.** Description, geographic coordinates, sampling date, water volume, number of replicates, depth and abbreviations.

**S2 Table. Pair of primers used to amplify the partial sequences of COI, 12SrRNA, and 16SrRNA genes. Sequences and references are shown for each marker.**

**S3 Table. Summary of reads, Amplicon Sequence Variants (ASVs) and Molecular Operational Taxonomic Units (MOTUs) statistics.** Based on eDNA analyses of both pilot and station sampling, values are shown after and before trimming steps.

**S4 Table. List of Amplicon Sequence Variants (ASVs) amplified using eDNA metabarcoding.** For both pilot and sampling stations included, the molecular marker, frequency, number of samples, length and taxonomic classification information are displayed for each ASVs.

**S5 Table. Biodiversity inventory for non-fish taxa in the Três Irmãos (TI) reservoir.** The inventory was built using previous literature, eDNA and GBIF information. For each taxon, the source of the record is indicated, as well as the reference, if needed.

**S6 Table**. **Biodiversity inventory for ichthyofauna in the Três Irmãos (TI) reservoir.** The inventory was built using previous literature, eDNA and Global Biodiversity Information Facility (GBIF) information. For each taxon, the source of the record is indicated, as well as the reference, if needed.

